# Impact of CRISPR/HDR-editing versus lentiviral transduction on long-term engraftment and clonal dynamics of HSPCs in rhesus macaques

**DOI:** 10.1101/2023.12.13.571396

**Authors:** Byung-Chul Lee, Ashley Gin, Chuanfeng Wu, Komudi Singh, Max Grice, Ryland Mortlock, Diana Abraham, Xing Fan, Yifan Zhou, Aisha AlJanahi, Uimook Choi, Suk See de Ravin, Taehoon Shin, Sogun Hong, Cynthia E. Dunbar

## Abstract

For precise genome editing via CRISPR/homology-directed repair (HDR), effective and safe editing of long-term engrafting hematopoietic stem cells (LT-HSCs) requires both sufficient HDR efficiency and protection of LT-HSC function and number. The impact of HDR on true LT-HSCs clonal dynamics in a relevant large animal model has not previously been studied. To track the HDR-edited cells, autologous rhesus macaque (RM) CD34^+^ cells were electroporated with the gRNA/Cas9 ribonucleoprotein (RNP) and HDR cassette barcode library structure and reinfused into RMs following myeloablation. For competitive model animals, fractionated CD34^+^ cells were transduced with a barcoded GFP-expressing lentiviral vector (LV) and electroporated via HDR machinery, respectively. *CD33* knockout (KO) neutrophils were prevalent early following engraftment and then rapidly decreased, resulting in less than 1% total editing efficiency. Interestingly, in competitive animals, a higher concentration of i53 mRNA result in a less steep reduction in *CD33* KO cells, presented a modest decrease in HDR rate (0.1-0.2%) and total indels (1.5-6.5%). In contrast, the drop off of LV-transduced GFP^+^ cells stabilized at 20% after 2 months. We next retrieved embedded barcodes and revealed that various clones contributed to early hematopoietic reconstitution, then after dominant clones appeared at steady state throughout the animals. In conclusion, CRISPR/HDR edited cells disappeared rapidly after the autologous transplantation in RM despite substantial gene editing outcome, whereas LV-transduced cells were relatively well maintained. Clonality of HDR-edited cells drastically shrank at early stage and then relied on several dominant clones, which can be mildly mitigated by the introduction of i53 mRNA.

## INTRODUCTION

The advent of CRISPR/Cas9 gene editing technologies has revolutionized the development of hematopoietic stem and progenitor cell (HSPC) gene therapies (*1*). Adapted from prokaryotic antiviral machinery, CRISPR technology relies on the introduction of RNA-guided DNA double stranded breaks (DSBs) by the Cas9 or related endonucleases and DSB repair by two evolutionarily conserved repair pathways: non-homologous end joining (NHEJ) or homology-directed repair (HDR) (*2, 3*). Error-prone NHEJ pathways result in small insertions or deletions (INDELs) at target sites, resulting in disruption of gene or regulatory functions, rather than mutation correction. In contrast, co-delivery of a homologous cassette can result in precise HDR-mediated genetic correction or gene insertion, with multiple potential therapeutic applications.

Despite the precision of HDR, it rapidly became apparent that the efficiency of HDR editing was generally much lower than NHEJ, particularly in HSPCs (*4, 5*). HSPCs also appear to be particularly susceptible to apoptosis resulting from activation of cellular pathways such as TP53 activation in response to DSBs, intracellular immunity in response to HDR cassette delivery via non-integrating viral vectors or large oligonucleotides, and toxicity related to the electroporation of editing components (*6–10*). In addition, recruitment of p53-binding protein 1 (53BP1) to DSB sites enhances NHEJ by inhibiting key HDR reactions. Inhibition of both TP53-mediated apoptosis and this pathway by i53, a ubiquitin variant inhibitor of 53BP1, has been shown to improve HDR efficiency and decrease HSPC toxicity (*11–14*). Design of guide RNAs favoring specific DNA repair mechanisms, optimizing the length of flanking arms in the HDR donor cassettes, improving component delivery, stimulating cell cycle progression, and expression of HDR cellular machinery components have also increased HDR efficiency in HSPCs (*13–17*).

Previous studies have predominantly utilized immunodeficient murine models to assess HDR editing efficiency and function of *in vivo* engrafting HSPCs, and several groups have recently reported significant progress towards the levels of HDR editing relevant for clinical utility (*12, 18, 19*). However, extrapolation of results from these xenograft models to long-term HSPC persistence and integrity in clinical applications is challenged by aspects of xenograft models, including abnormal microenvironment, lack of completed differentiation for some lineages, competition from residual murine HSPCs, short follow-up based on murine lifespan, and impossibility of assessing most safety parameters. Non-human primates (NHP) such as rhesus macaques (RMs) overcome many of these hurdles, closely modeling human lifespan and size, immune milieu, HSPC dynamics, marrow microenvironment, and safety parameters (*20–22*). NHP models have been used previously to assess and optimize HSPC viral gene addition therapies and NHEJ editing, predicting clinical trial results (*21, 23, 24*).

Polyclonality is the desired goal following HSPC gene therapies and serves as a quantitative measure of the impact of *ex vivo* manipulations on stem cell numbers and function (*13, 25*). Oligoclonality and clonal expansions are associated with an increased risk of malignant transformation and other outcomes related to HSPC proliferative stress and dysfunction (*26*). Assessment of clonality following transplantation is thus highly relevant to understanding the impact of HDR on HSPC function long term. We have previously reported that inclusion of a high diversity barcode library within integrated viral vectors in a RM model allows sensitive and quantitative tracking of clonal output from transduced HSPCs (*27, 28*). Inclusion of genetic barcodes within HDR cassettes or other clonal tracking techniques have also been applied to study clonality in murine xenograft models (*13*).

We now utilized the RM autologous transplantation model to assess the impact of CRISPR/HDR editing on long-term multilineage HSPC engraftment and clonal dynamics at a single-cell level by inclusion of a barcode library within the HDR cassette. Given the robust preclinical and clinical experience with HSPC lentiviral gene addition therapies, and ongoing clinical trials for hemoglobinopathies using both gene therapy approaches, we have directly compared the impact of each methodology on HSPC long term engraftment and clonality. Understanding the comparative long-term clonal dynamics of CRISPR/HDR-edited versus lentiviral-edited HSPCs will provide key insights into the predicted comparative clinical safety and efficacy of these approaches.

## RESULTS

### Optimization of CRISPR/HDR gene editing at the *CD33* locus

To study the impact of CRISPR/HDR editing on RM HSPCs, we chose to target the rhesus *CD33* gene. We previously reported no discernable changes in HSPC dynamics or differentiation following CRISPR/NHEJ editing and knockout of function at this locus in the RM autologous transplantation model (*24*). Using the same *CD33* gRNA previously utilized for NHEJ (*24*), median INDEL frequency was 42% in primary RM CD34^+^ cells, however HDR knock-in was very low at less than 1% (Fig. S1A). According to recent publications (*29, 30*), microhomology-mediated end joining (MMEJ)-biased gRNAs favor HDR outcomes, therefore, we used to LINDEL algorithm to design ten additional *CD33* gRNAs predicted to favor MMEJ (Table S1; Fig. S1B) and tested them for HDR efficiency using an ssODN cassette and Cas9/gRNA ribonucleoprotein complexes (RNP) in primary RM CD34^+^ cells. Targeted deep sequencing revealed that HDR efficiency was highest for gRNA #5 targeting exon 2 (Fig. 1A, B). To confirm functional inactivation of CD33, the CD33 positive human monocytic leukemia cell line MOLM-14 was electroporated with gRNA #5 and CD33 expression was assessed by flow cytometry. Homozygous KO and loss of *CD33* expression in a substantial fraction of cells was observed (Fig. S1C). We co-electroporated i53 mRNA along with the ssODN HDR cassette and RNP into CD34^+^ cells to improve HSPC HDR gene editing, based on prior experience in human HSPCs (*11*). A mean 26.3% increase in HDR was achieved in the presence of i53 mRNA (Fig. 1C). These conditions and gRNA were taken forward into *in vivo* macaque studies.

**Figure 1.**
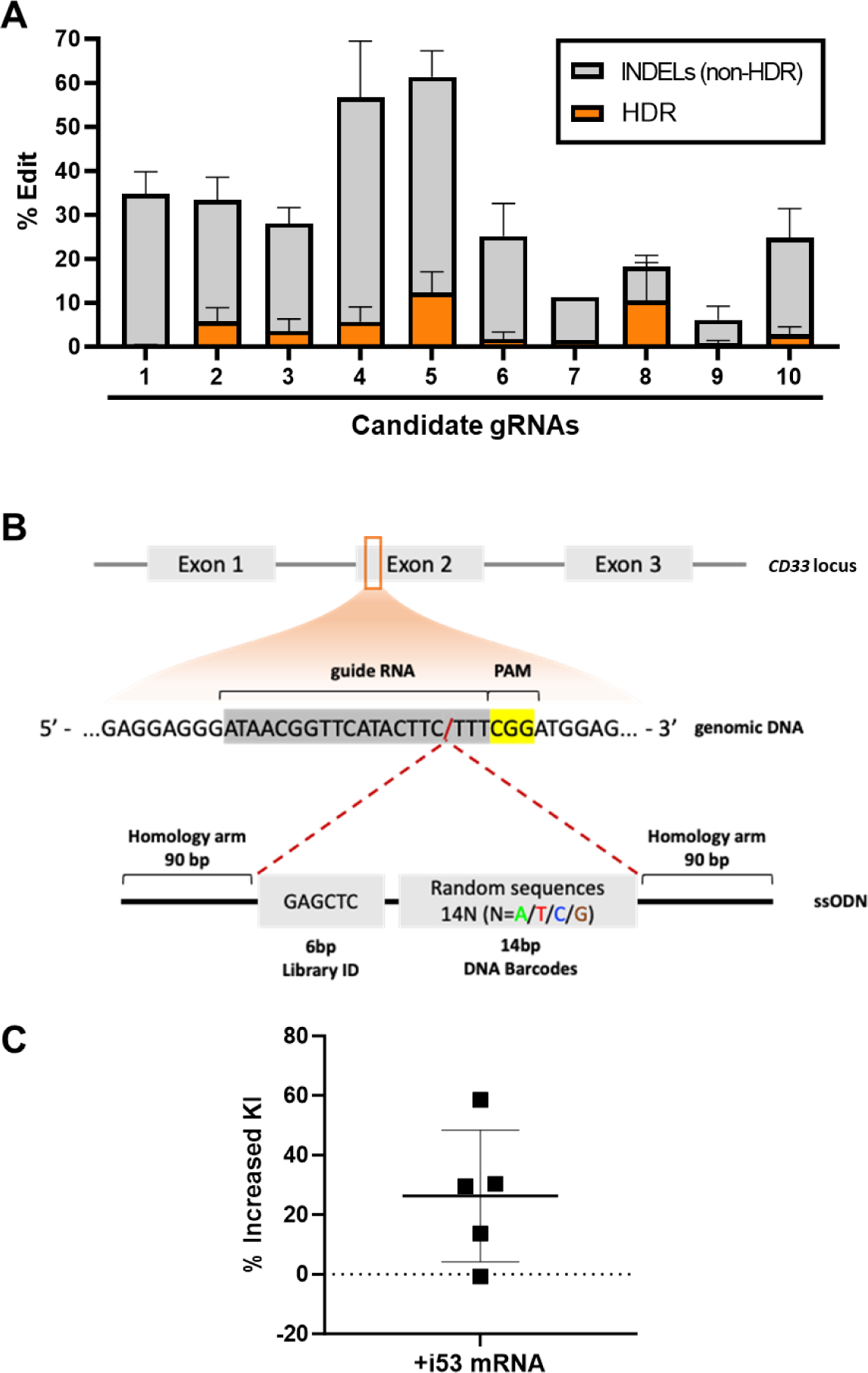
*In vitro* optimization of CRISPR/HDR editing at *CD33* genetic locus. **(A)** Mutation frequencies of candidate gRNAs in primary macaque CD34^+^ cells as assessed by high throughput targeted deep sequencing 5 days after electroporation. Non-HDR indel frequency and HDR-mediated cassette insertion are indicated in grey and orange respectively. **(B)** Schematogram of CRISPR/HDR gene editing with barcoded cassette insertion at the rhesus macaque *CD33* locus. **(C)** Percent improvement in HDR-mediated knock-in efficiency via inclusion of i53 mRNA (1 μg/ml) during editing of in CD34^+^ primary rhesus macaque HSPCs as assessd by deep sequencing of celle maintained *in vitro* for 5 days following electroporation of editing components (n=5 independent experiments).

### Transplantation of *CD33* CRISPR/HDR barcoded cells

As diagramed in Figure 2A, CD34-enriched HSPCs from Animal #1 were cultured for 24-48 hours then electroporated with Cas9/*CD33* gRNA#5 RNP, a high diversity barcoded single stranded ODN HDR donor cassette, and i53 mRNA (1 μg/ml). 24 hours following electroporation, 6.4 million CD34^+^ HSPCs/kg were reinfused into the autologous macaque following myeloablative total body irradiation (TBI) conditioning, after removal of aliquots for analysis of the infusion product (IP) (Table 1). INDEL (non-HDR) allelic frequency was 61.3% and HDR 15.4% (Fig. 2B). To confirm disruption of the *CD33* locus via NHEJ INDELs or HDR, IP CD34^+^ cells were placed into myeloid differentiation culture for 14 days and CD33 expression was assessed by flow cytometry. Non-edited controls cells were 43% CD33^+^ in contrast to only 4.3% CD33^+^ following editing, confirming substantial homozygous *CD33* locus disruption (Fig. 2C). Genotyping of individual CFU plated from the IP confirmed a high frequency of both homozygous and heterozygous *CD33* disruption, with only 12.5% unedited CFU (Fig. 2D). Furthermore, the ratios of NHEJ vs. HDR alleles was approximately 3:1, reflecting the similar proportions of both alleles observed in the IP (Fig. 2B and D).

**Figure 2.**
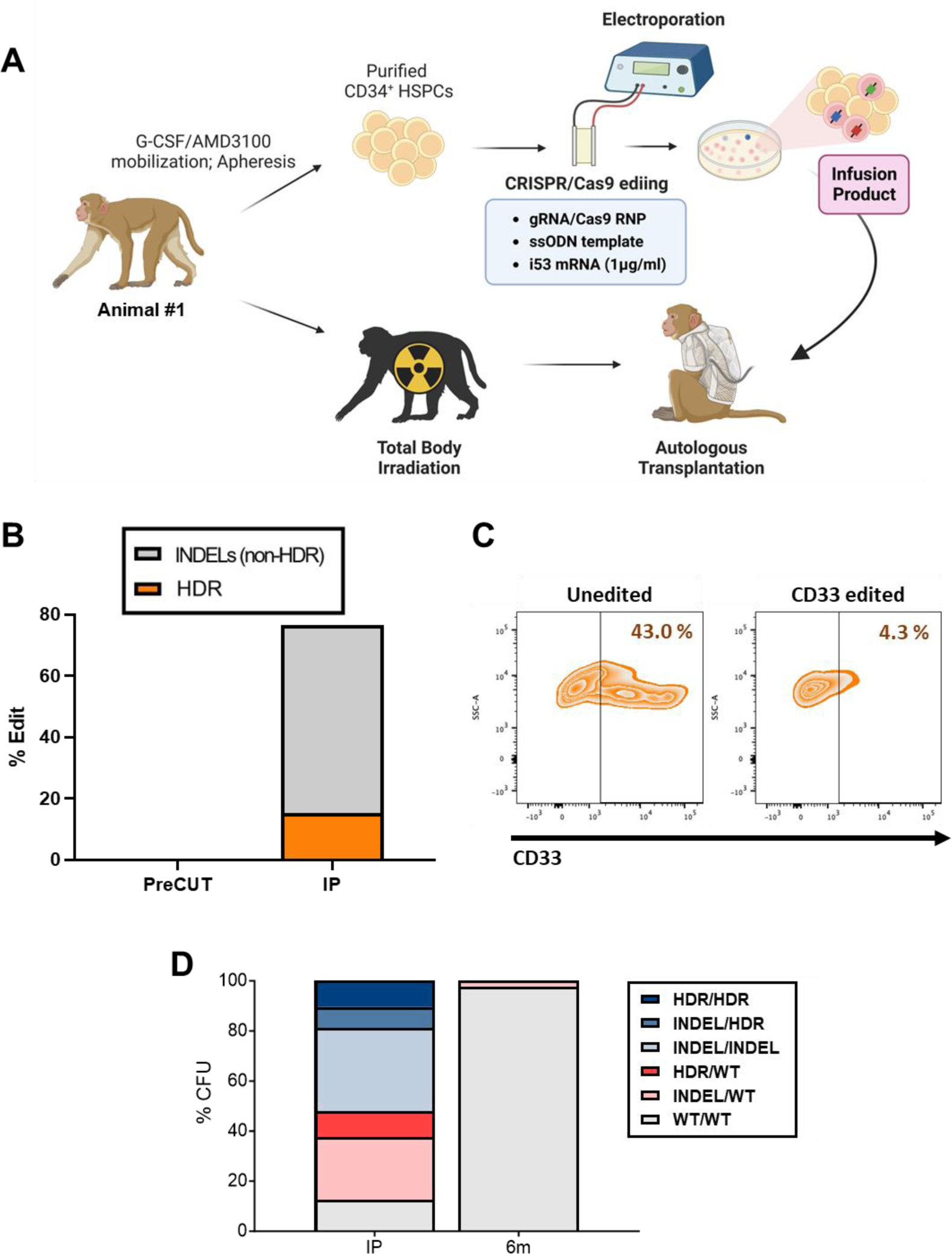
Rhesus macaque CRISPR/HDR gene editing autologous transplantation model. **(A)** Schematic of CD34^+^ HSPC cell collection, *CD33* target site gene editing and autologous transplantation performed for Animal #1. **(B)** Mutation frequency at target site *CD33* in the infusion product (IP) assessed via targeted deep sequencing 1 day following electroporation of the editing components. Non-HDR indel frequency and HDR-mediated cassette insertion are indicated in grey and orange respectively. **(C)** Evaluation of CD33 expression by flow cytometry in un-edited versus an aliquot of *CD33*-edited cells from Animal #1’s infusion product following *in vitro* culture under myeloid differentiation conditions for 14 days. **(D)** Genotyping of individual CFU grown from the *CD33-*edited infusion product (IP) or from bone marrow CD34^+^ HSPCs obtained from the animal 6 months post-transplantation. Each CFU was classified based on all possible combinations of HDR versus INDEL versus WT alleles.

**Table 1.** Summary of animals and transplantation parameters.

|  | Animal #1 | Animal #2 | Animal #3 |
| --- | --- | --- | --- |
| Age at transplantation | 12yr 3m | 4yr 9m | 5yr |
| Gender | Female | Female | Female |
| Body weight (kg) | 5.7 | 4.7 | 6 |
| CD34<sup>+</sup> cells infused (x 10e6) | 36 | 11.05 (CRISPR) | 10.46 (CRISPR) |
|  |  | 11.69 (Lenti) | 21.21 (Lenti) |
| Total CD34<sup>+</sup> cells/kg (x 10e6) | 6.4 | 4.9 | 5.28 |
| Viability of infused cells (%) | 79.5 | 75.3 (CRISPR) | 83.5 (CRISPR) |
|  |  | 98.2 (Lenti) | 87.7 (Lenti) |

Following transplantation, we monitored engraftment via an assessment of complete blood counts (CBC) over time as compared to previously transplanted animals conditioned with the same dose of TBI and receiving autologous HSPCs NHEJ-edited at the *CD33* locus or lentivirally transduced (Fig. S2). The timing of recovery of neutrophils and lymphocytes were similar to these prior comparator animals, and recovery of red blood cell production appeared somewhat delayed but within the expected range. However, platelet recovery was delayed compared to comparator animals, with animal 1 requiring 5 platelet transfusions, the last 19 days post-infusion.

To track *in vivo* repopulation with CRISPR/HDR barcoded HSPCs, we examined circulating granulocytes (Gr) over time, a short-lived population that best reflects ongoing HSPC output (*31*). Targeted deep sequencing of Grs at the time of initial engraftment 2 weeks post-transplantation showed INDEL and HDR knock-in (KI) allelic frequencies in Grs to be 46.8% and 3%, respectively (Fig. 3A). However, levels of both total INDELs and HDR alleles subsequently rapidly decreased, dropping to 0.68% and 0.008% respectively at longest follow-up of 24 months post-transplantation. Homozygous CD33^-^ KO Grs as detected by flow cytometry also rapidly decreased, reaching baseline levels by 3 months post-transplantation (Fig. 3B). The absence of homozygous-edited CD34^+^ cells was validated by marrow single colony CFU genotyping performed 6 months post-transplantation (Fig. 2D). We also determined editing levels in purified B cells, T cells, and NK cells, as well as in marrow CD34^+^ cells (Fig. S3A). The pattern in B cells and NK cells matched total INDELs and HDR levels in Gr, with rapid decreases following initial higher engraftment levels. Edited T cells appeared later and at low levels, as expected for new T cell output following thymic damage resulting from TBI conditioning.

**Figure 3.**
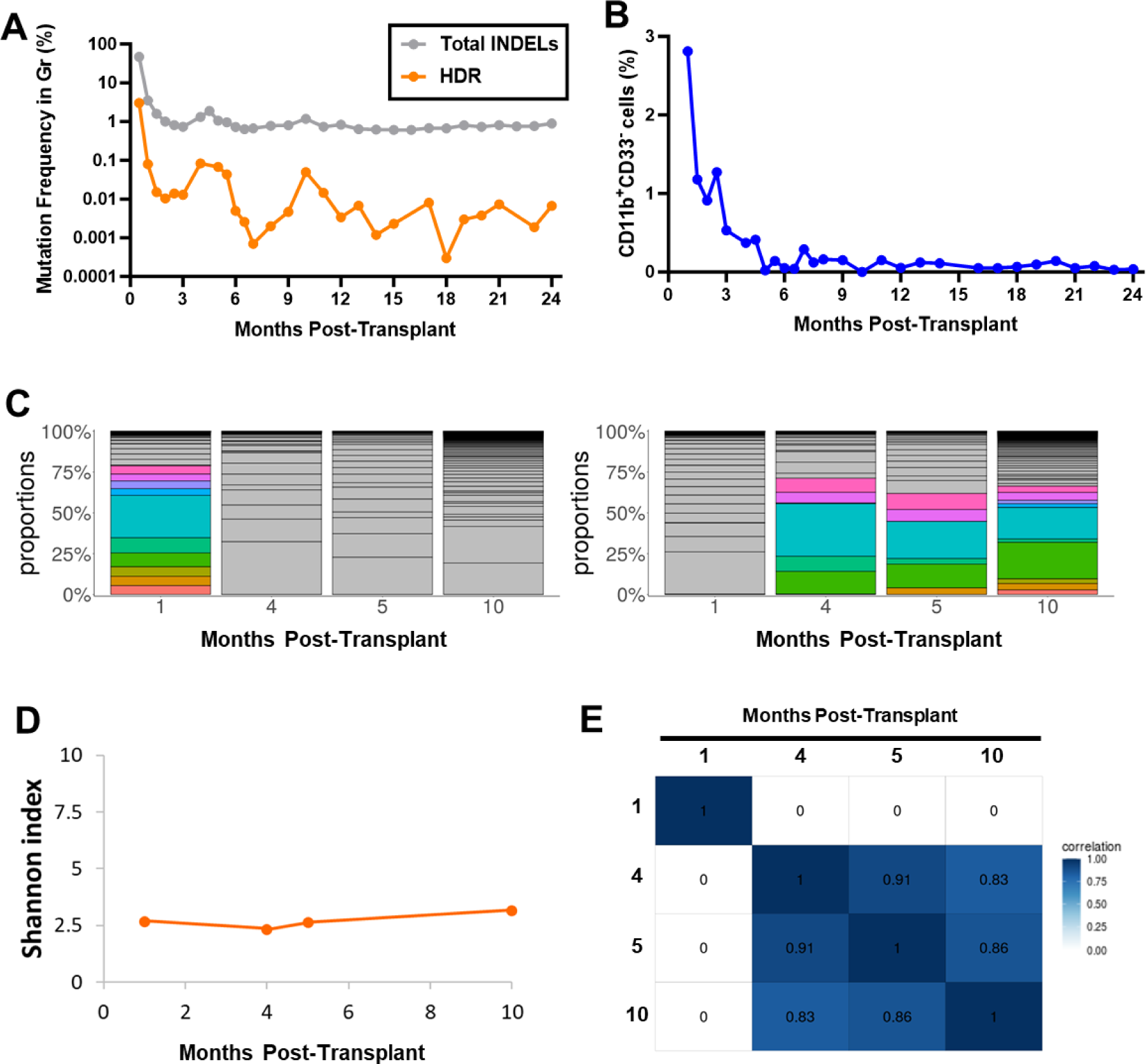
Editing frequencies and clonality in Animal #1. **(A)** Total INDEL and HDR editing allelic percentages over time post-transplantation in peripheral blood granulocytes as assessed by targeted deep sequencing of the *CD33* locus. Total INDELs and HDR-mediated knock-in indicated with grey and orange lines respectively. **(B)** Longitudinal monitoring of the loss of CD33 expression on CD11b^+^ peripheral blood granulocytes by flow cytometry. **(C-E)** Analysis of the clonality of CRISPR/HDR-mediated inserted barcodes retrieved from peripheral blood granulocytes via PCR, deep sequencing, and application of a custom analysis pipeline at indicated time points in months post-transplantation. **(C)** Stacked plots show the percentage contribution of the 10 largest contributing clones in PB granulocytes at a designated time points mapped over the other time points. Each of the clones at the index time point is shown as separate colors in the stack, with the same clone designated by the same color at the other time points if present. Contributions from other clones are shown via grey scale. The contributions from the top 10 contributing clones at 1 month (left) and 10 months (right) post-transplantation are shown. **(D)** Shannon diversity of CRISPR/HDR barcodes over time. The Shannon diversity index encompasses both the number of clones and the evenness of their distribution. **(E)** Clonal relationships at a population level as depicted by Pearson correlation coefficients comparing pairwise fractional contributions from all valid barcodes for the samples at indicated time points with color intensity representing *r* values for each comparison.

### Clonal dynamics of engrafted HSPCs

We performed deep sequencing to retrieve barcodes inserted appropriately via HDR into the *CD33* locus, and utilized the Barseq pipeline for HDR barcode extraction, modified to fit our experimental conditions (*13, 32*). We identified a sampling threshold by performing duplicate barcode retrieval from Gr samples of various DNA amounts, achieving > 96% detection in both duplicates with 500ng of DNA, thus this DNA amount was used for all barcode retrieval (Fig. S4). Samples with sufficient DNA for barcode retrieval from 1, 4, 5, and 10 months post-transplantation were analyzed. Clones detected as contributing at 1m had become undetectable at later time points, replaced by another set of clones found at 4, 5 and 10m, mirroring the pattern we have previously reported following lentiviral barcoding, with independent contributions from short-term versus long-term engrafting HSPCs (Fig. 3C) (*27, 28*). The top 10 dominant clones at the later time points contributed up to 75% of all barcoded contributions, suggesting oligoclonal HDR-edited hematopoiesis (Fig. 3C and D), confirmed by Shannon diversity indexes. This clonal pattern was then stable over time between 4 and 10m, with a very high degree of clonal similarity between the timepoints (Fig. 3E). In sum, very few HDR-edited long-term repopulating clones survived *in vivo*, not surprising given the very low overall level of HDR editing.

### Impact of lentiviral transduction versus HDR editing on dynamics of genetically modified HSPCs in a competitive transplantation model

To directly compare the impact of CRISPR HDR editing versus lentiviral gene addition, on the function of HSPCs *in vivo*, we performed competitive transplantation experiments in two macaques (Fig. 4A, Table 1, Animals #2 and #3). Both approaches require *ex vivo* culture and manipulation of HSPCs and are being developed therapeutically for several human disorders, with years of safety and efficacy data available for lentiviral transduction. For each animal, CD34-enriched HSPCs cells were divided into 2 equal aliquots. One aliquot was transduced with lentiviral vector (LV) expressing CopGFP and containing the high diversity barcode library we have previously utilized to study hematopoiesis and aspects of gene addition therapies (*28, 33, 34*). LV transduction efficiency of CD34^+^ cells was 35-40% as assessed by flow cytometry for GFP^+^ expression in the IP (Fig. 4B and S5A). The second aliquot was electroporated with the *CD33* RNP and barcoded ssODN HDR cassette, as well as a higher concentration of i53 mRNA compared to Animal #1 in order to potentially further improve HDR editing efficiency and HSPC viability (*12, 35–37*). The editing efficiencies for total INDELs and HDR in the CD34^+^ HSPCs were similar to those achieved in Animal #1 (Fig. 4C), with both homozygous and heterozygous editing in individual CFU, and only approximately 10% unedited CFU (Fig. 4D). The editing outcomes in the IP overall were reflected in the colonies. After myeloid differentiation culture, CD33 expression in CD11b^+^ cells decreased by 34.7% and 29.3%, respectively, compared to LV-transduced cells (Fig. 4E and S5B). Plating efficiency of CFU was lower for edited versus transduced CD34^+^ cells (Fig. S5C). The two aliquots were combined and reinfused into the autologous animal following TBI conditioning (Fig. 4A). Transduction and transplantation parameters are given in Table 1 for Animals #2 and #3. Both animals engrafted and recovered all blood lineages promptly, in contrast to the delayed platelet recovery observed in Animal #1 (HDR only) (Fig. S2).

**Figure 4.**
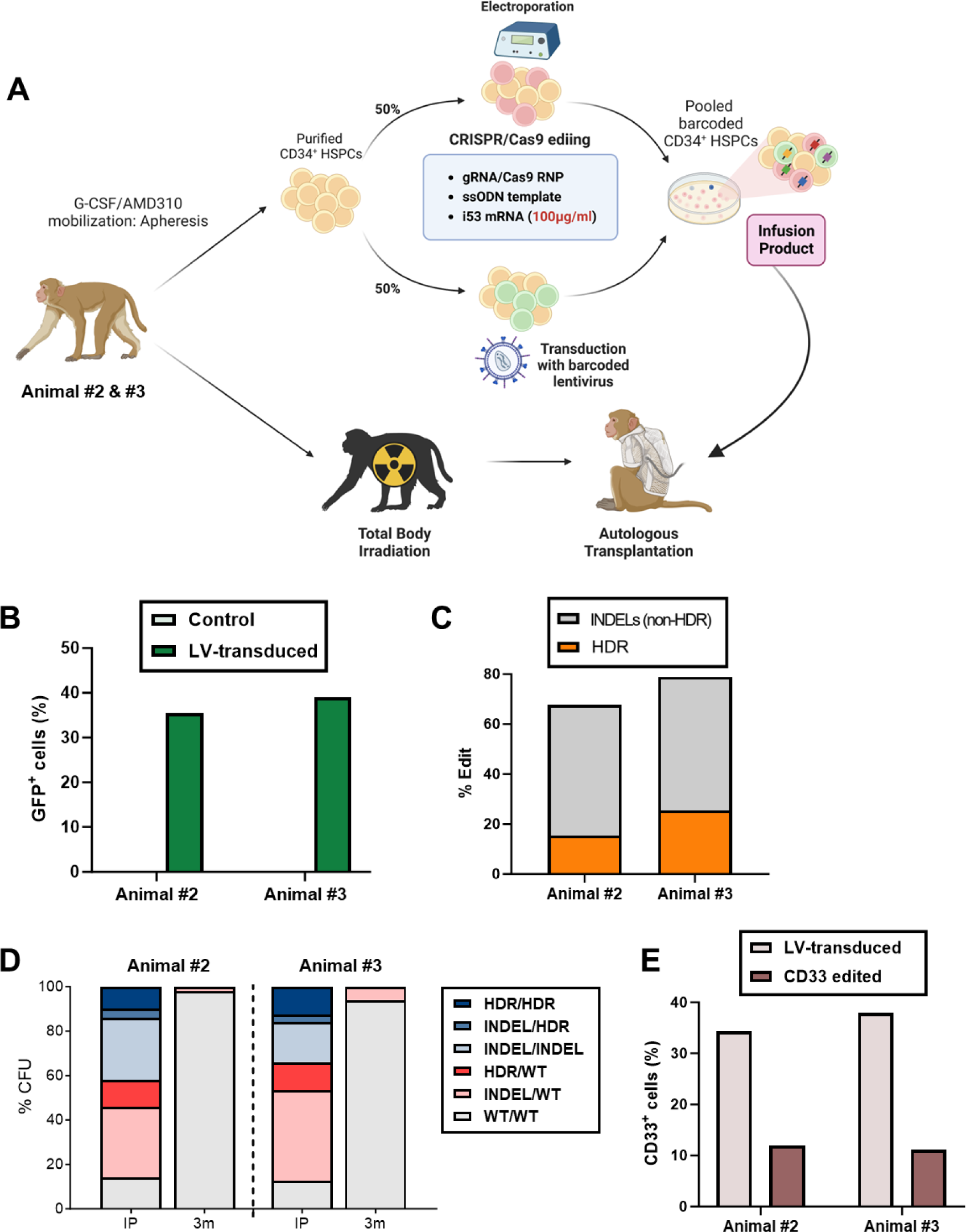
Competitive rhesus macaque autologous transplantation model comparing CRISPR editing to lentiviral transduction in Animals #2 and #3. **(A)** Schematic of CD34^+^ HSPC cell collection, *CD33* target site gene barcoded editing versus barcoded lentiviral transduction, followed by competitive autologous transplantation as performed in Animals #2 and #3. **(B)** GFP expression by flow cytometry in aliquots of the lentivirally-transduced CD34^+^ HPSCs from Animals #2 and #3 assessed 3 days following transduction. **(C)** Mutation frequency at the *CD33* target site in aliquots of the CRISPR edited CD34^+^ HSPCs from Animals #2 and #3 assessed by targeted deep sequencing 1 day following electroporation. Non-HDR indel frequency and HDR-mediated cassette insertion are indicated in grey and orange respectively. **(D)** Genotyping of individual CFU grown from the *CD33-*edited infusion product (IP) or from bone marrow CD34^+^ HSPCs obtained 3 months post-transplantation shown for Animals #2 (left) and #3 (right). Each CFU was classified based on all possible combinations of HDR versus INDEL versus WT alleles. **(E)** Aliquots of lentivirally transduced or CRISPR edited cells were cultured for 14 days in myeloid differentation conditions, and CD33 expression was evaluated by flow cytometry.

We assessed the fraction of GFP^+^ Grs originating from the LV-transduced arm over time (Fig. 5A), with long-term marking stabilizing at approximately 15-22% in both animals after initial engraftment that was slightly higher, particularly in Animal #2, a pattern similar to our prior reports, suggesting somewhat better transduction of short-term versus long-term engrafting cells, and at levels as expected or higher, given that only half the starting HSPCs were LV-transduced. Both HDR-edited and total INDEL alleles in Grs from Animals #2 and #3 were detected at frequencies higher than in Animal #1, with a 5 to 15-fold higher levels of total INDELs and 20 to 100-fold increases in HDR alleles, despite only half the cells having been exposed to editing in Animals #2 and #3 compared to Animal #1, and competition from the LV-transduced aliquot, suggesting that the higher concentration of i53 had positive impact on HSPC function (Fig. 5B). However, HDR editing levels long-term were still less than 1%. Long-term engrafting cells with homozygous editing were not sustained over time, as evidenced by complete loss of CD33^-^ myeloid cells (Fig. 5C) and lack of homozygously-edited CFU by 3 months post-transplantation in both animals (Fig. 4D). Analysis of GFP^+^, HDR-edited and total INDELs in other lineages and marrow CD34^+^ cells showed the same pattern as Gr, other than the expected delay in production of edited T cells (Fig. S3B-D).

**Figure 5.**
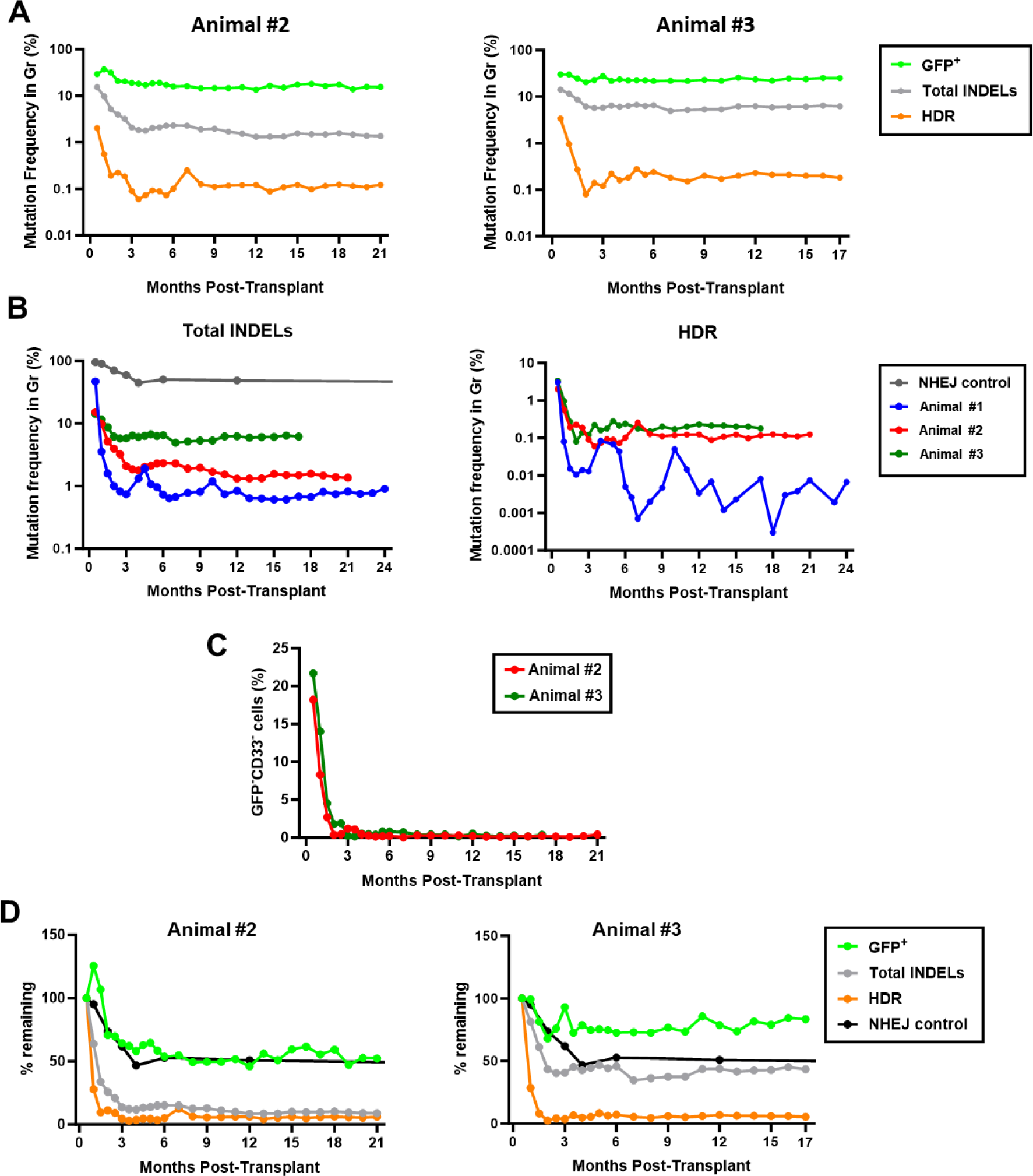
Tracking of HDR, INDELS and GFP over time following autologous transplantation. **(A)** Comparison of lentiviral transduction levels (as assessed by %GFP expression, shown in green) to HDR-edited (orange) or INDEL (grey) alleles over time in peripheral blood granulocytes from Animals #2 (left) and #3 (right). **(B)** Comparison of INDELs in granulocytes over time post-transplantation from all 3 current animals and a previous animal receiving cells edited at the *CD33* locus without an HDR cassette (left) and HDR for all three current animals (right). **(C)** Fraction of CD33^-^GFP^-^CD11b^+^ granulocytes over time post-transplantation monitored by flow cytometry for Animals #2 and #3. **(D)** Plot of the percent GFP, HDR or INDEL levels remaining in granulocytes over time, as compared to the earliest levels measured at 2 weeks post-transplantation, in Animals #2 and #3, and in comparison to the NHEJ only animal. Calculated by dividing the percentage present at week two by the percentage at each later time point and expressing as a percentage.

To compare the impact on short versus long term engrafting cells with LV versus CRISPR/HDR editing, we normalized Gr marking to the level observed at 2 weeks and plotted the % marking remaining over time for GFP (LV transduction), HDR and total INDELs (Fig. 5D). LV-transduced long-term HSPC output was sustained at 50-75% of the 2 week maximum, while HDR dropped to less than 5% of maximum. For total INDELs, animal 3 maintained just less than 50% of maximum long-term, similar to prior NHEJ animals (Fig. 5D), however Animal #2’s LT-HSPC total INDELs dropped to less than 10% of maximum. The results strongly suggest impairment or loss of long-term HSPCs during HDR-editing, not only for HDR edited cells, but also potentially for NHEJ edited cells exposed to HDR editing conditions, and in stark contrast to much better-preserved output from LV-transduced HSPCs.

### Clonality following CRISPR/HDR editing versus LV-transduction of HSPCs

To address clonal dynamics of engrafted CRISPR/HDR-edited and LV-transduced cells, retrieved barcodes were analyzed. The LV barcode pattern was highly polyclonal and diverse, with 9,000-almost 16,000 clones detected in total from each animal, and 3,000-4,000 individual stable LT-HSPCs contributing at later time points, with even the largest clones never contributing more than 1-2% (Fig. 6 and S6). As previously observed, short-term repopulating (ST)-HSPC lineage-restricted clones contributed immediately after engraftment and then disappeared, replaced by stable multipotent LT-HPSCs (Fig. S7). Likely due to the higher concentration of i53 mRNA during CRISPR/HDR editing, we retrieved more stable and consistent HDR-barcoded clones from Animals #2 and #3 over time compared to Animal #1. However, in contrast to hematopoiesis from LV-transduced HSPCs, output from HDR-edited cells was much more oligoclonal long-term, with several large clones representing more than 5-30% each of HDR-edited Gr long term, with many fewer clones detected compared to LV clones at later time points, and lower diversity (Fig. 6 and S6). This is not surprising given the much lower level of cells being produced from HDR versus LV engineered cells long term. It is however important to note that the HDR-edited clonality pattern overall was similar to the LV pattern, with replacement of early lineage-restricted ST-HSPC clones by stable multilineage LT-HSPC (Fig. S7).

**Figure 6.**
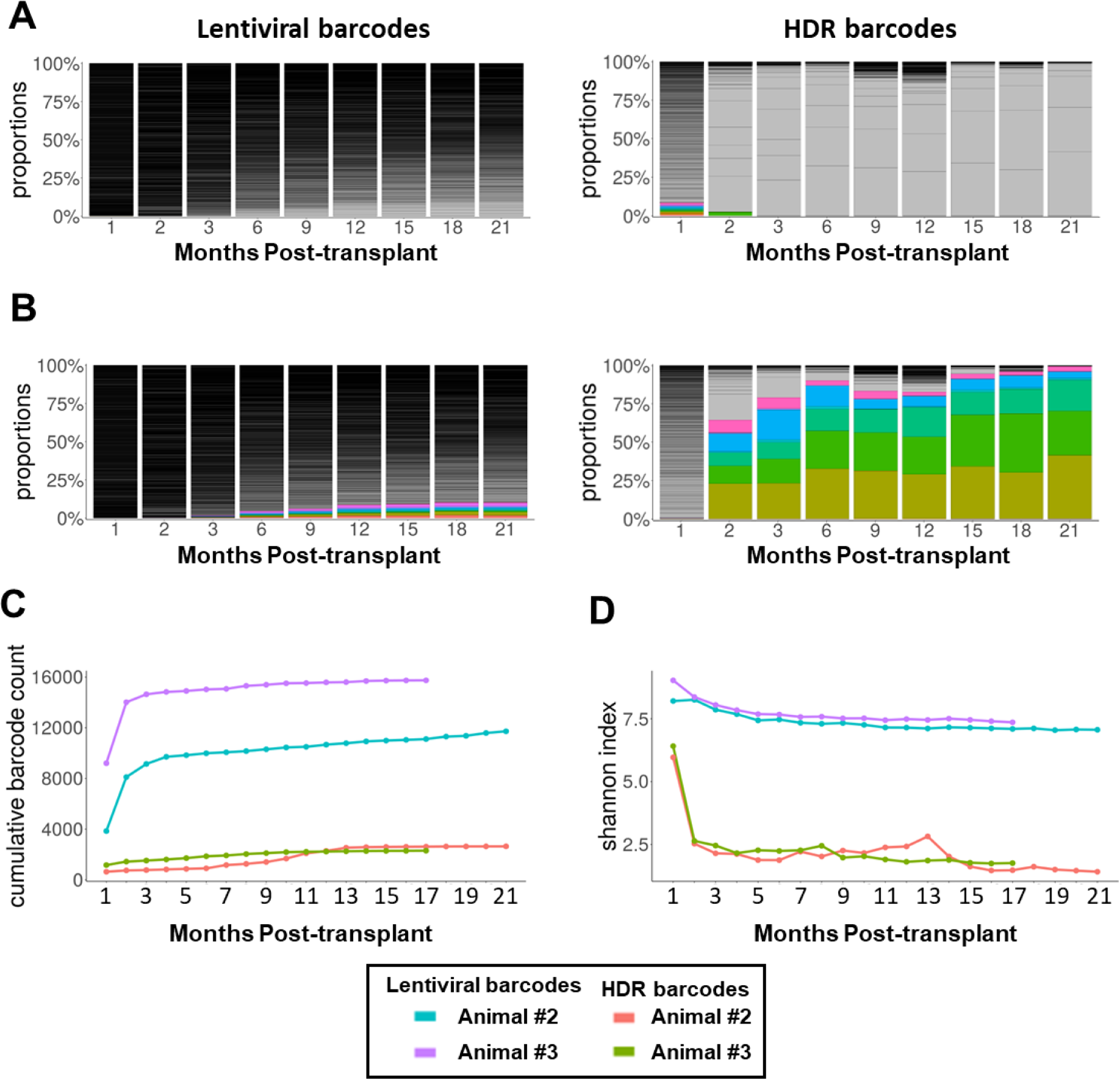
Clonal contribution analysis of RM *Lentiviral-CRISPR*/HDR competitive animals. Inserted barcodes from both lentiviral (LV) arm and CRISPR/HDR arm were retrieved from PB granulocytes via BAR Seq. and custom analysis pipeline over time. **(A-B)** The stacked bar plots for percentage of the largest top 10 contributing clones in PB Grs of Animal #2 over time. Each of the clones is shown as separate colors in the stack. Contributions from the other clones are shown as a single solid color. The fraction of contributing clones at 1 month **(A)** and 21 months **(B)** post-transplantation were presented. Left: Lentiviral barcodes, Right: HDR barcodes. **(C)** Cumulative number of independent barcoded clones originating from each HSPC arm in Animals #2 and #3 over time. **(D)** Shannon diversity for each arm in Animals #2 and #3. The Shannon diversity index encompasses both the number of clones and the evenness of their distribution.

### Off-target editing

We identified predicted off target editing sites for the *CD33* gRNA utilized via an *in silico* algorithm (see Methods) (Fig. S8A). Targeted deep sequencing of the top 11 predicted sites was performed on granulocytes from pre-transplantation and 3 months post-transplantation blood in all 3 animals. No off target editing at any site was detected in blood cells (Fig. S8B-D).

## DISCUSSION

In the present study, we investigated long-term *in vivo* engraftment and clonal dynamics of transplanted CRISPR/Cas9 HDR-edited autologous RM HSPCs, a highly relevant and predictive preclinical model (*20*). We directly compared the impact of HDR editing on HSPC engraftment and clonality to LV gene addition, an HPSC engineering modality with over a decade of clinical experience and extensive prior NHP comparator studies. The current rapid expansion of HSPC editing modalities into human clinical trials, and the active development of both LV gene addition and editing approaches for disorders such as sickle cell anemia and severe immunodeficiencies make these comparisons highly relevant. Tracking engineered HSPCs over up to two years post-transplantation, we found that only a small fraction of CRISPR/HDR edited cells engrafted and contributed to hematopoiesis long-term, presumably due to functional damage to true long-term HSPCs during the HDR gene editing process. Although more potent inhibition of p53 activation improved HDR-edited cell engraftment, absolute levels and clonal diversity long term remained far inferior to LV-transduced HSPC compared directly in a competitive model. The recent announcement of prolonged cytopenias and lack of substantial engraftment with HDR-corrected alleles in the first patient enrolled in a trial attempting to correct the sickle cell disease (SCD) mutation, resulting discontinuation of the program by the sponsor, emphasizes the importance of predictive large animal models for new modalities entering clinical development (*38*).

Almost all research regarding *in vivo* efficacy of gene insertion or correction via HDR has been carried out via administration of edited human HSPCs to immunodeficient mice. Several recent xenograft studies have documented several orders of magnitude better long-term engraftment with HDR-modified human HSPCs than achieved in our RM study (*4, 35, 37, 39–45*), and such data was used to support moving forward into clinical trials, including the recently halted SCD study. Prior comparative clonal studies have suggested that NHP HSPCs engrafting in immunodeficient mice do not represent those engrafting in autologous NHPs (*46, 47*). Behavior of human or NHP HSPCs in the highly abnormal xenogeneic microenvironment may be very different from that in a native homologous marrow microenvironment, due non-cross-reactive cytokines and HSPC-matrix interactions (*48*). Cues driving HSC self-renewal or differentiation may be lacking or non-physiologic, given ongoing production of mature cells from residual murine HSPCs and limited production of red cells, platelets and other mature human blood cells (*49, 50*).

In our study, both short- and particularly long-term engraftment with HDR-edited as well as NHEJ-edited cells was far lower than that documented in the infusion product. A high editing efficiency achieved *in vitro* prior to infusion may not directly correlate with *in vivo* engraftment and persistence. Macaques in both the current study and the prior published NHP study documented a decline not only in HDR knock in alleles but also in NHEJ and MMEJ alleles (*37*). In addition, delayed platelet recovery occurred in Animal #1 in our study and in an animal enrolled in the prior study, both receiving only cells exposed to HDR editing, also suggesting general toxicity from the *in vitro* HDR-editing processes. These observations indicate that direct or indirect damage can impact on all HSPCs exposed to the *ex vivo* HDR procedure, rather than solely HSPCs with successfully-HDR edited alleles, perhaps due to interferons and other stress signals released from cells during the editing process.

We optimized the experimental condition to improve HDR-mediated integration by selection of gRNAs favoring MMEJ repair pathways and inclusion of an mRNA encoding an inhibitor of p53 to block alternate NHEJ repair and apoptosis resulting from double stranded DNA breaks (*11, 12, 29, 30, 36, 51*), resulting in an increase in HDR knock-in rates at the *CD33* target gene site from less than 1% to as high as 26%, and overall NHEJ/MMEJ/HDR editing efficiency of 80% in macaque CD34^+^ HSPCs prior to transplantation. These improvements did not result in sustained high levels of HDR editing *in vivo* in the macaques. TP53 activation is higher with inclusion of donor HDR templates as compared to gRNA/endonucleases alone (*52*). However, recent studies in xenografted mice found that p53 inhibition increased the fraction of HDR-edited engrafted HSPCs (*13, 51*), in contrast to our results in primates, further suggesting differences between cells engrafting in xenografts and in a native environment. According to a recent study, susceptibility to p53-mediated toxicity can vary depending on the DNA repair mechanisms (*53*), with more p53-related toxicity during NHEJ than HDR, suggesting other mechanisms of HDR-related toxicity.

We have demonstrated that cells edited through the HDR machinery in all three macaques exhibited profoundly oligoclonal hematopoiesis in myeloid and lymphoid lineages, particularly long-term, in comparison to highly polyclonal infusion products, suggesting loss of LT-HSCs with the editing procedure. Clonal tracking in xenograft models have also shown loss of clonal diversity over time, however less profound than in our macaques, and in part restored via p53 inhibition along exposure to an adenoviral protein improving cell cycle progress and HDR efficiency (*13*). Of note, another group reported most contributions from a few highly dominant clones in primary xenograft recipients, despite detection of many additional smaller clones, however in secondary recipients, very few clones were detected, suggesting that LT-HSPCs successfully undergoing and surviving HDR were rare (*54*). Besides the implications for slow hematopoietic recovery and low efficacy with loss of HDR-corrected cells, oligoclonality is also worrisome for later development of neoplasia due to proliferative stress on a small number of remaining stem cells.

While HSPC LV gene addition has been explore for over almost two decades in multiple clinical trials, with clear clinical successes in several serious disorders and resultant in regulatory approvals, theoretical and now documented concerns regarding genotoxic leukemias resulting from semi-random vector insertions have accelerated interest in targetable gene editing therapies. In addition, LV gene addition approaches are not applicable to many autosomal dominant disorders, in contrast to mutation correction via HDR, however both approaches are being pursued in several disorders including SCD and immunodeficiencies, thus direct comparisons are desirable. While in our primate model, CRISPR/HDR was far less promising regarding efficiency and clonality, one direct comparison in a xenograft model was more encouraging. Brault and coworkers reported even better engraftment with HDR gene-corrected compared to LV-transduced HSPCs from patients with X-SCID, and improved *in vitro* NK development (*55*). Besides the differences detailed above regarding autologous macaque versus xenograft models, the use of X-SCID as compared to normal target HSPCs results in selection for corrected lymphoid progeny and can overcome efficiency constraints. No clonal analyses were performed in these studies.

Our study focused solely on editing at the *CD33*, chosen for its clinical relevance and ability to make comparisons to animals already transplanted with CRISPR/NHEJ edited HSPCs (*24, 56*). Additionally, we utilized ssODN instead of AAV6 or non-integrating lentiviral vectors as homologous donor templates, based literature suggesting that while HDR efficiency *in vitro* is higher with AAV6 template delivery (*57, 58*), long-term *in vivo* engraftment even in xenograft models was significantly higher with ssODN delivery (*45*). An assessment of the impact of AAV6 on long-term engraftment in large animal models is desirable.

Much progress has been made utilizing gene correction via HDR to develop treatments for many serious diseases, targeting cell types in tissues other than HSPCs. However, the current primate study highlights some of the unique barriers and risks associated with HDR targeting HSPCs. Approaches including base editing or prime editing that allow targeted mutation correction without inducing DSBs are very exciting, with encouraging murine and xenograft model data (*59–61*). We look forward to further investigation of these approaches in non-human primate models.

## MATERIALS AND METHODS

### Rhesus macaque HSPC autologous transplantation

All animal experiments were conducted according to protocols approved by the National Heart, Lung, and Blood Institute (NHLBI) Animal Care and Use Committee, following institutional and Department of Health and Human Services guidelines. RM HSPC mobilization, collection, autologous transplantation and post-transplantation supportive care were performed as described (*24, 62*). In brief, HSPCs were mobilized by subcutaneous administration of granulocyte-colony stimulating factor (G-CSF) (Amgen, Thousand Oaks, CA) 15 μg/kg/day for 6 days, and AMD3100 (Sigma-Aldrich, St. Louis, MO) 1 mg/kg on days 5 and 6, followed by apheresis using a Fenwal CS-3000 cell separator on days 5 and 6. CD34^+^ cells were enriched via immunoselection using anti-mouse IgM microbeads (Miltenyi Biotec, Bergisch Gladbach, Germany) and the anti-CD34 hybridoma clone 12.8 (a murine IgM anti-human CD34 cross-reactive with RM CD34), derived at the Fred Hutchinson Cancer Center, Seattle, WA and not commercially available. Following myeloablative total body irradiation (TBI) at a dose of 4.5Gy rads daily for two days, autologous CD34^+^ cells were infused following TBI on the second day.

For Animal #1, all infused CD34^+^ cells were CRISPR/HDR-edited as detailed below. For Animals #2 and #3, collected CD34^+^ cells were split into two equal fractions. One was CRISPR/HDR-edited and the other was lentivirally transduced. Following completion of editing or transduction, the cell fractions were collected, pooled, and infused together into the autologous recipient.

### Gene editing of hematopoietic cells

To form *CD33* gRNA/Cas9 RNP editing complexes, *CD33* chemically modified guide RNAs (6.25 μg/10^6^ cells, Synthego, Redwood City, CA) and Cas9 protein (12.5 μg/10^6^ cells, PNA Bio, Thousand Oaks, CA) were mixed for 10 minutes at room temperature. gRNA sequences targeting the *CD33* locus are given in Table S1. Screening of gRNA editing efficiency was performed on CD34^+^ cells and on the MOLM-14 cell line for analysis of CD33 expression. For targeted insertion of a barcode via HDR, 200 nucleotides (nt) length single-stranded oligo deoxynucleotides (ssODN) were synthesized (Ultramer^TM^ DNA oligonucleotides, Integrated DNA Technologies (IDT), Coralville, IA), consisting of two 90 nt homology arms flanking the *CD33* target site surrounding a 6 nt barcode library ID and a 14 nt high diversity random barcode (Fig. 1B and Table S1). p53 inhibitor i53 mRNAs (Table S1) were synthesized using the mMessage mMachine T7 Transcription Kit (Invitrogen, Carlsbad, CA) and bulk i53 mRNA for Animals #2 and #3 was obtained from CELLSCRIPT, LLC (Madison, WI).

CD34^+^ cells collected from each animal on the two days of apheresis were placed in culture for 48 hours in X-VIVO^TM^ 10 (Lonza, Walkersville, MD) supplemented with 1% human serum albumin (HSA) (Baxter, Deerfield, IL) and human recombinant cytokines stem cell factor (SCF) 100 ng/mL, FMS-like tyrosine kinase 3 ligand (FLT3L) 100 ng/mL, and thrombopoietin (TPO) 100 ng/mL; (all from PeproTech, Rocky Hill, NJ). The next day, CD34^+^ cells were resuspended in Opti-MEM (Lonza) at a concentration of 3-5×10^6^ cells in a total volume of 750 μl. The cell suspension was mixed with ssODNs (1 ug/1×10^6^ cells), i53 mRNA (1 μg/ml) for Animal #1, 100 μg/ml for Animals #2 and #3), and RNPs were electroporated with a single pulse of 400V for 5 msec using the BTX ECM 830 Square Wave Electroporation System (BTX/Harvard Apparatus, Holliston, MA). Electroporated CD34^+^ cells were placed back into the same culture media overnight before differentiation cultures, colony-forming unit assays, or reinfusion into the autologous RMs.

### Lentiviral transduction of rhesus macaque HSPCs

Barcoded lentiviral vectors (LV) containing a 6 bp library identifier followed by a 27 bp random barcode and expressing the CopGFP transgene were produced, and barcode diversity of the vector library measured as previously described (*28, 63*). Fractionated CD34^+^ cells from Animals #2 and #3 were cultured overnight on RetroNectin-coated plates (Takara, T100B, Mountain View, CA) in X-VIVO^TM^ 10 (Lonza) supplemented with 1% HSA (Baxter) and with 100 ng/ml each of human FLT3L, SCF, and TPO (PeproTech) and transduced the next day with barcode-containing LV at a multiplicity of infection (MOI) of 25 in the presence of 4 mg/ml protamine sulfate (Sigma) (*28*). 24 hours later, transduced CD34^+^ cells were pooled with gene edited cells and reinfused into irradiated autologous RMs 2 and 3.

### Assessment of editing efficiencies

Genomic DNA (gDNA) was extracted from edited RM CD34^+^ HSPC using the DNeasy Blood & Tissue kit (QIAGEN, Germantown, MD) and gRNA target regions were amplified by PCR with primers given in Tables S1 and S2. Editing frequencies in samples collected following transplantation were assessed via deep sequencing of PCR amplified libraries of the *CD33* gRNA target region (Table S1) on an Illumina Miseq (Illumina, San Diego, CA) via 300bp paired end reads and a median sequencing depth of 100,000 reads/gRNA target site. To define single nucleotide polymorphisms (SNPs), consensus sequences for the *CD33* target locus were determined for each animal via sequencing of pre-transplantation blood samples from each individual RM. Sequencing reads were demultiplexed and processed using MiSeq Reporter (Illumina). Alignment of amplicon sequences to an unedited consensus sequence was performed using CRISPResso2 (http://crispresso2.pinellolab.org/) and the number of reads quantified for each unique sequence. To quantify all the editing events, total editing efficiency was calculated as a percentage of (the number of reads with indels)/(the number of total reads). For assessment of HDR-mediated insertion at the target region as a percentage of (the number of reads with the 20 nt barcode insertion)/(the number of total reads).

### Barcode retrieval and analysis

500ng of DNA from each sample was used for both LV and HDR barcode retrieval via 30-cycle PCR. Primers are given in Table S2. To allow for multiplex sequencing of pooled samples, the forward primer contained a unique i5 index for each sample paired with a universal reverse primer. For HDR barcode retrieval, the forward primer contained flanking sequences around the target *CD33* site and the 6bp barcode ID to amplify only HDR inserted barcoded alleles (Table S2). For LV barcode retrieval, primers flanking the barcode region within the vector were used (Table S2) (*28*). After gel purification of the PCR products, 40-80 multiplexed samples were pooled for sequencing on an Illumina NovaSeq 6000 system to recover the barcodes.

Barcoded extraction was conducted using the Bar-Seq analysis pipeline with some modifications (*13*). Fastq files were processed with TagDust (v.2.33) to extract the inserted barcode from each sample by first aligning to the HDR or LV barcode library ID, and especially upstream and downstream *CD33* flanking sequences or vector sequences in the HDR and LV arms respectively. Analyses and visualizations were performed using R (Foundation for Statistical Computing) and Prism (GraphPad Software, San Diego, CA). Custom R code has been described and is available on GitHub at https://github.com/dunbarlabNIH/barcodetrackR (*64*).

### Blood and marrow collection and lineage purification

Rhesus macaque peripheral blood (PB) was separated via centrifugation over Lymphocyte Separation Medium (MP Biomedicals, Santa Ana, CA). PBMNC (peripheral blood mononuclear cells) and granulocyte (Gr) fractions were treated with ACK lysing buffer (Quality Biological, Gaithersburg, MD) to remove red blood cells. Cells for lineage analyses were stained with the following antibodies: CD3-BV786, CD20-APC-cy7, CD16-APC (BD Biosciences, San Jose, CA), CD14-pacific blue (Thermo Fisher Scientific, San Jose, CA), CD159a-PE-cy7 (NKG2a, Beckman Coulter), CD11b-FITC (BioLegend, San Diego, CA), and CD33-PE (Miltenyi Biotec), and flow cytometric analysis and/or sorting performed on a FACSARIA-II (BD Biosciences). Data were analyzed using FlowJo software (Tree-star Inc., Ashland, OR).

10 ml of bone marrow was aspirated from the posterior iliac crests or ischial tuberosities 6 months (Animal #1) or 3 months (Animals #2 and #3) post-transplantation, mononuclear cells isolated and CD34-enriched via column immunoadsorption as described above.

### Myeloid differentiation

Post-editing, aliquots of CD34^+^ cells were cultured in Iscove’s Modified Dulbecco’s Medium (IMDM) (Gibco-BRL, Grand Island, NY) supplemented with 10% fetal bovine serum (FBS), and human SCF (100 ng/ml), FLT3 (100 ng/ml), TPO (50 ng/ml), interleukin 6 (IL-6) (50 ng/ml), granulocyte-macrophage colony-stimulating factor (GM-CSF) (100 ng/ml), and rhesus interleukin 3 (IL-3)(10 ng/ml) (all Peprotech) for 2 to 3 weeks at 37°C in a humidified atmosphere with 5% CO_2_. The cells were maintained at a concentration of 5-10×10^5^/ml with medium changes every 3 to 4 days. The expression of CD11b and CD33 were analyzed by flow cytometry as above.

### Colony forming unit (CFU) assays

CD34^+^ cells from the infusion product or purified from bone marrow mononuclear cells post-transplantation were plated in MethoCult H4435 Enriched media (STEMCELL Technologies, Vancouver, Canada) in 35-mm dishes at a concentration of 1000 cells/ml and incubated at 37°C and 5% CO_2_. After 14 days of culture, individual colonies were evaluated and plucked for DNA extraction.

### Off-target analyses

Off-target sites were selected as previously described (*65*), using an in-house Python script modified for the rhesus macaque genome from the published prediction algorithm (https://zlab.bio/guide-design-resources). Each potential off-target site that has 3 or less mismatches to the gRNA were scored and among them, the top 11 predicted sites were selected for design of flanking PCR primers (Table S2). Illumina sequencing of the products was then performed.

### Statistical analysis

All statistical analyses were conducted using GraphPad Prism 9. Unpaired Student’s t test was performed for pairwise comparisons and one-way analysis of variance (ANOVA) with Bonferroni’s multiple comparison post-test for three or more groups, as indicated.

## Acknowledgments

The authors would like to acknowledge Theresa Engels, Justin Golomb, Allen Krouse, and N. Seth Linde for excellent animal care, Aylin Bonifacino for technical assistance, and the NHLBI Genomics and Flow Cytometry Cores.

## Author contributions

B-C.L. and C.E.D. conceptualized the study. B-C.L., C.W., and C.E.D. designed the experiments. B-C.L., A.G., M.G., D.A., F.X., Y.Z, T.S., and S.G.H. collected the data. B-C.L., A.G., U.C., S.S.R., and C.W., analyzed the data. B-C.L., A.G., K.S., M.G., R.M., and A.A. performed computational analysis. B-C.L., A.G., and C.E.D. wrote and edited the manuscript.

## Data and materials availability

Custom Python code for RM targeted deep sequencing is available at https://github.com/shint3/CH_crispr_analysis.git/.

