## supplemental materials for "Impact of CRISPR/HDR-editing versus lentiviral transduction on long-term engraftment and clonal dynamics of HSPCs in rhesus macaques"

**Contents:**

Supplemental Figures (Figure S1-S8)
Supplemental Tables (Table S1-S2)

**Supplemental Figures**

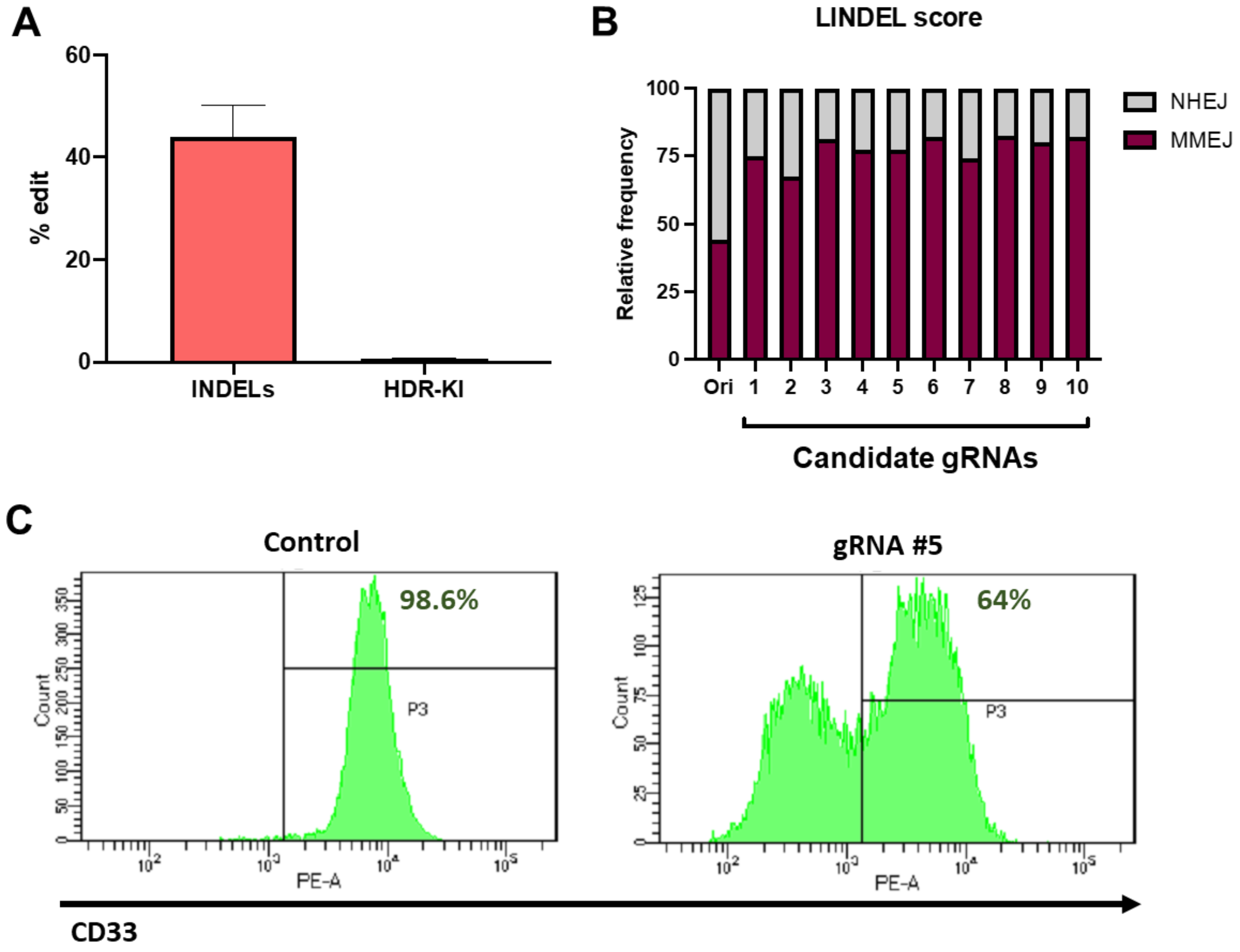


**Figure S1. gRNA selection and *in vitro* validation of CRISPR/HDR editing outcomes. (A)** Mutation frequencies of previously established chemically modified *CD33* gRNA in primary macaque CD34$^+$ cells by high throughput targeted deep sequencing 5 days after electroporation. Total indel frequency indicated in left side and HDR-mediated knock-ins (KIs) in right side. **(B)** *In silico* prediction of gene editing outcome of candidate *CD33* gRNAs via LINDEL scoring system. **(C)** RNP complexes including gRNA #5 showing exact homology to the human *CD33* target sequence were introduct to the CD33-expressing human myeloid cancer MOLM-14 cell line. Inhibition of CD33 expression was analyzed by flow cytometric analysis 5 days after electroporation.

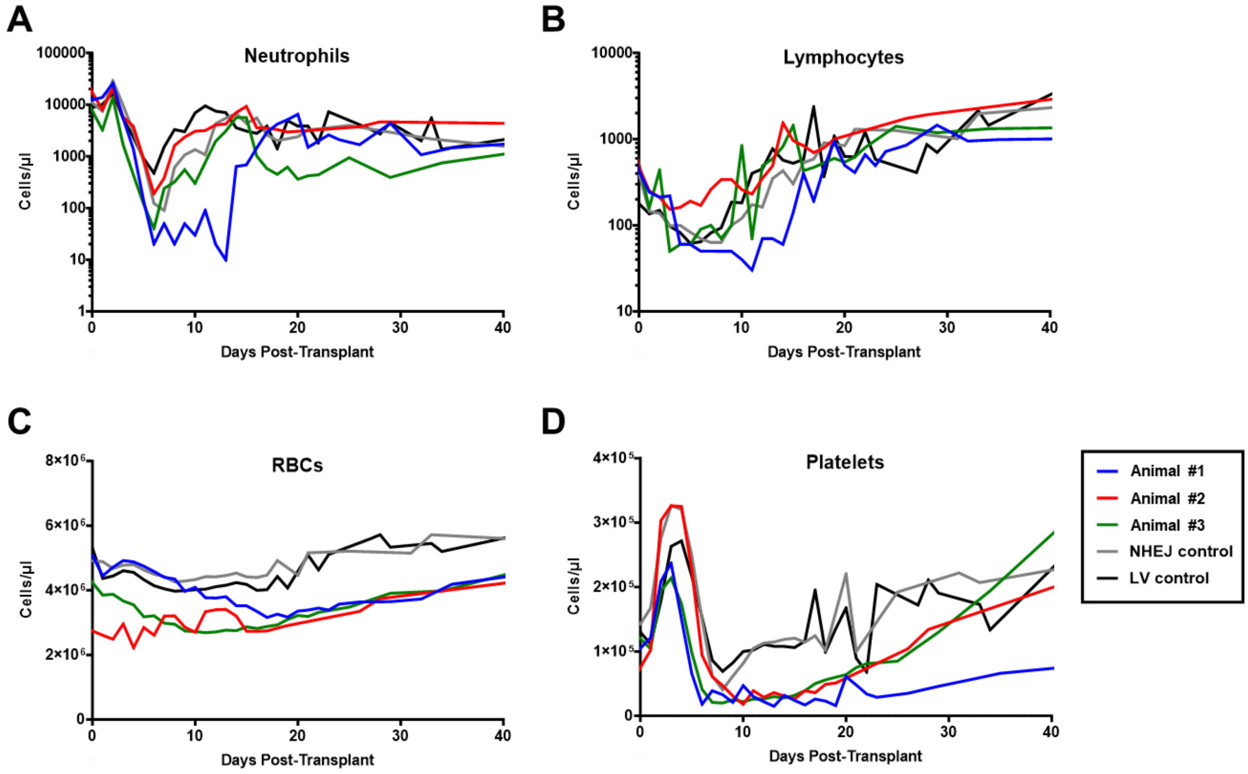


**Figure S2. Complete blood counts (CBCs) from animals for the first 40 days following autologous transplantation with CRISPR/HDR (Animal #1) or *Lentiviral-CRISPR*/HDR (Animals #2 and #3) HSPCs. (A)** Absolute neutrophil counts, **(B)** Lymphocyte counts, **(C)** Red blood cell counts (RBCs), and **(D)** Platelet counts. Each value is plotted over time and shown in comparison to a CRISPR/NHEJ control animal transplanted with cells edited at *CD33* locus and to mean values of lentivirally (LV) transduced control animals (n=4).

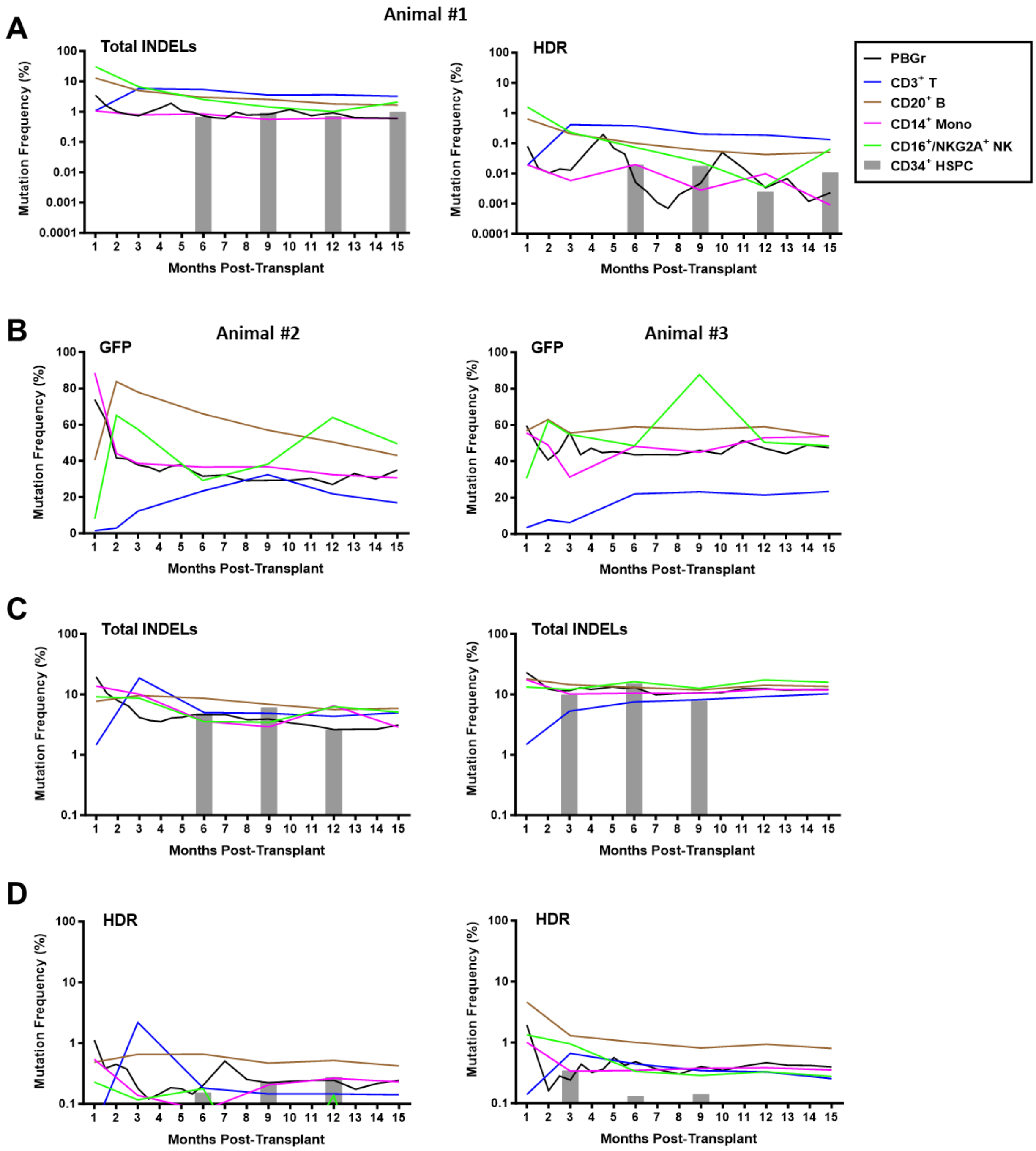


**Figure S3. *CD33* gene editing and lentiviral transduction efficiencies following autologous transplantation with CRISPR/HDR (Animal #1) or *Lentiviral-CRISPR/HDR* (Animal #2 and #3) HSPCs. (A)** *CD33* target site mutation frequencies by high throughput deep sequencing shown over time for purified peripheral blood (PB) T cells, B cells, monocytes and mature CD16$^{+}$ natural killer (NK) cells, and from CD34$^+$ HSPCs purified from the bone marrow (BM) analyzed for Animal #1. Total indels (left), CRISPR/HDR-mediated targeted knock-in (KI) (right). **(B)** Percentage of GFP$^+$ cells over time in PB purified granulocytes, T cells, B cells, monocytes, and mature CD16$^+$ NK cells over time. Left: Animal #2, Right: Animal #3. **(C-D)** *CD33* target site mutation frequencies by high throughput deep sequencing shown over time for purified PB T cells, B cells, monocytes and mature CD16$^+$ natural killer (NK) cells, and CD34$^+$ HSPCs from rhesus BM. Total indels **(C)** and CRISPR/HDR-mediated KI **(D)** are shown. Left: Animal #2, Right: Animal #3.

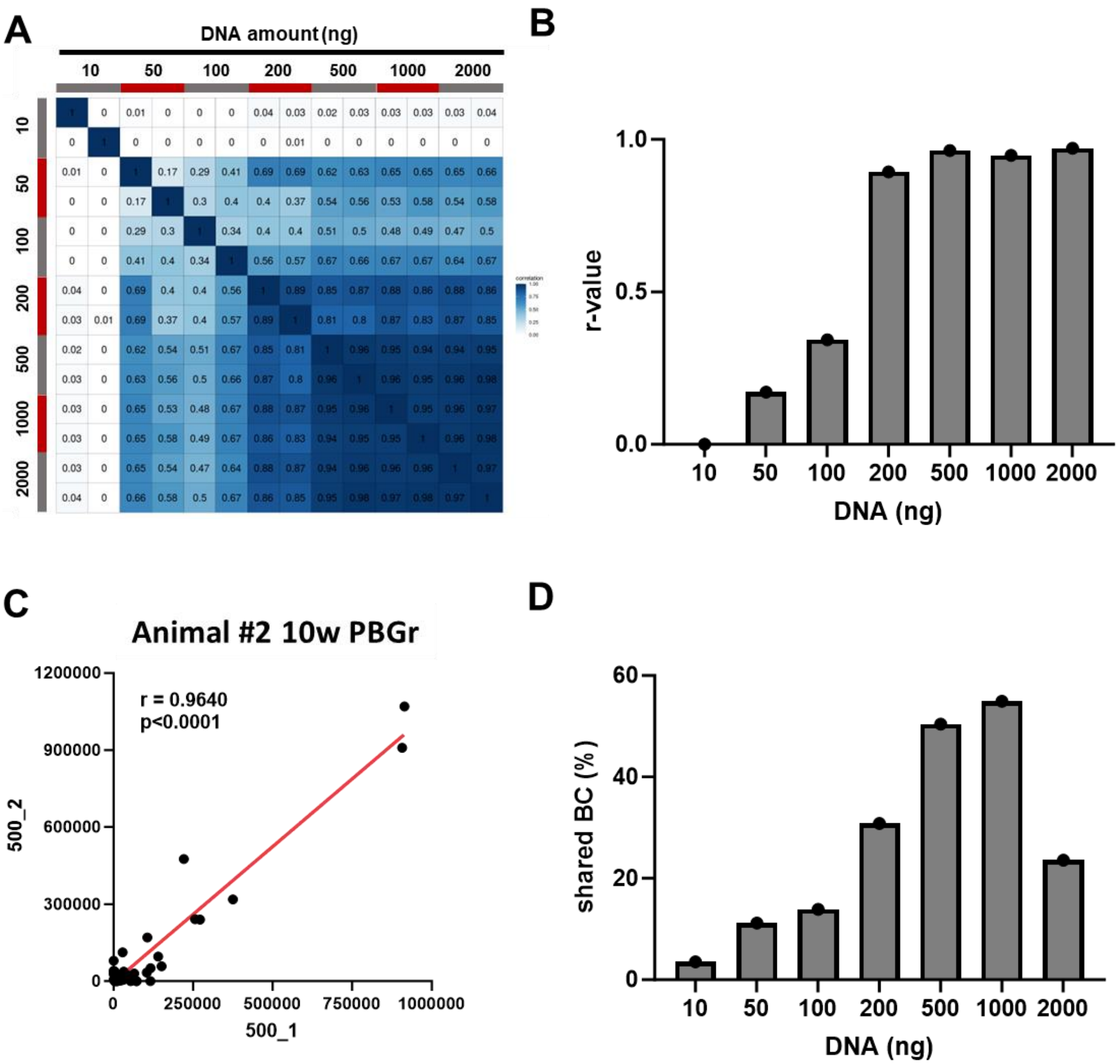


**Figure S4. Optimization of barcode retrieval.** A range of duplicate genomic DNA amounts obtained from Animal #2 granulocytes purified from the peripheral blood 10 weeks post transplantation were processed for barcode retrieval optimization. **(A)** Clonal relationships at a population level as depicted by Pearson correlation coefficients comparing pairwise fractional contributions from all valid barcodes for the duplicated samples. *r*-values are presented in each box. **(B)** *r*-values of each duplicated sample at designated DNA amounts are shown in the bar graph. **(C)** Detailed comparison of individual barcodes from duplicates of 500 ng of DNA. **(D)** Percentage of overlapping unique barcodes from various DNA amounts with those retrieved from 500 ng.

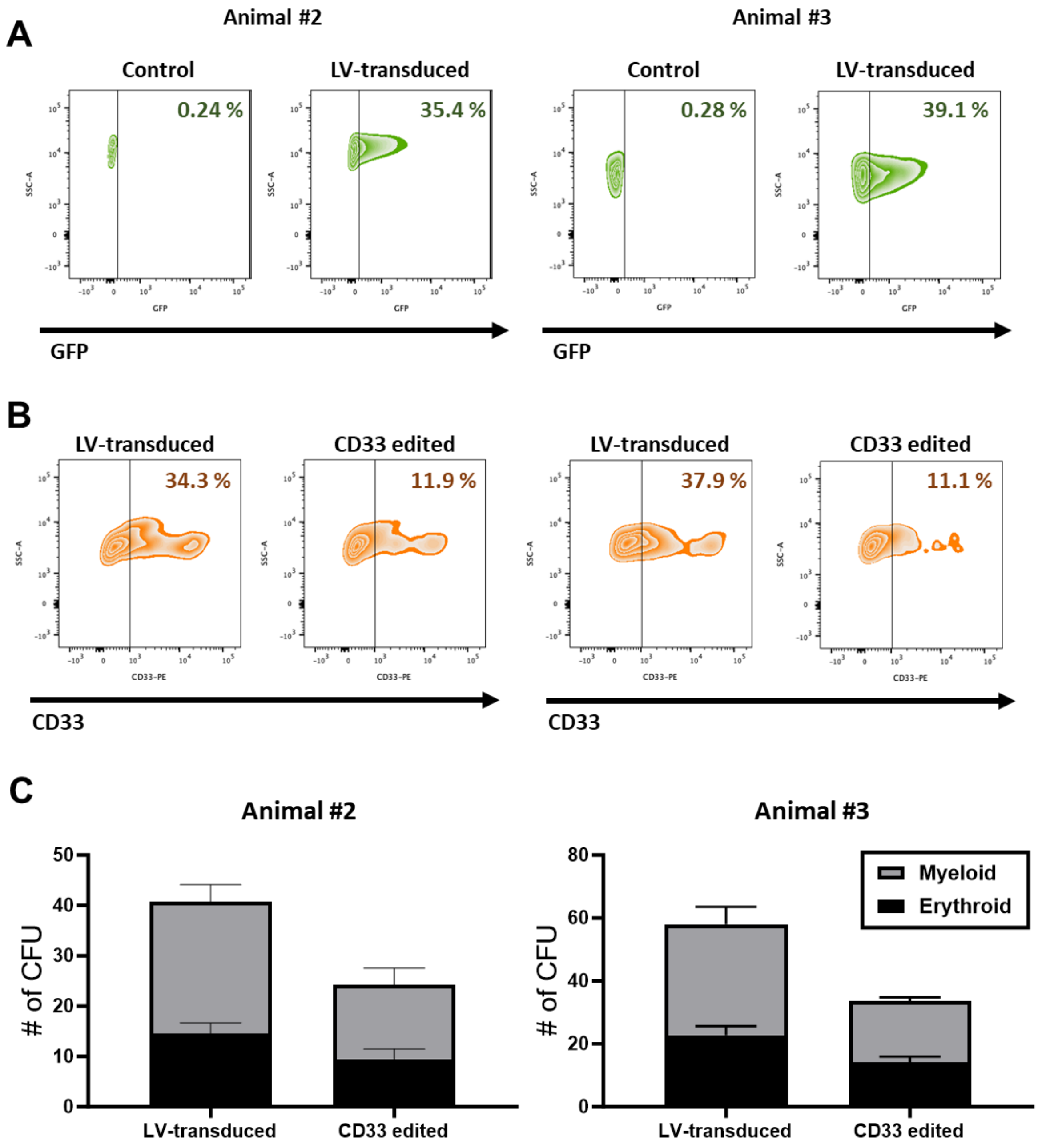


**Figure S5.** ***In vitro* validation of gene editing and LV transduction. (A)** Representative plot images of lentiviral GFP expression. GFP expression level from lentivirally transduced CD34$^+$ RM HSPCs of Animal #2 (left) and Animal #3 (right) were determined by flow cytometric analysis 3 days following transduction and *in vitro* culture. **(B)** Representative plot images of CD33 expression. The CD34$^+$ HSPCs were differentiated under myeloid conditions for 14 days *in vitro*

and CD33 expression was evaluated by flow cytometric analysis. The lentivirally transduced cells were used as a control comparison. Left: Animal #2, Right: Animal #3. **(C)** The number of colonies/plate from lentivirally (LV) transduced cells and CRISPR/HDR edited cells, seeded from the infusion product. Left: Animal #2, Right: Animal #3.

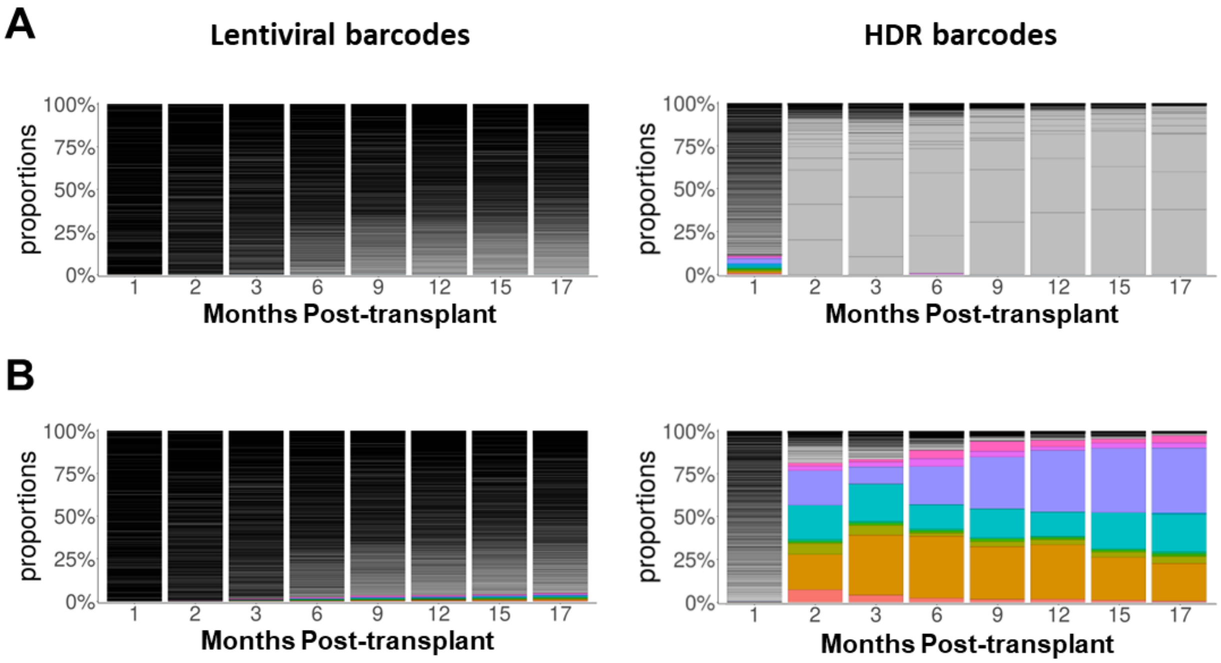


**Figure S6. Clonal contribution analyses of RM *Lentiviral-CRISPR/HDR* competitive Animal #3.** Inserted barcodes from both LV arm and CRISPR/HDR arm were retrieved from PB granulocytes (Grs) via BAR Seq. and application of a custom analysis pipeline. The stacked bar plots show the percentage of the 10 largest contributing clones in PB Grs of Animal #3 over time shown as individual colors in the stacked plot. Contributions from other clones are shown in grey scale. The fraction of contributing clones at 1 month **(A)** and 17 month **(B)** post-transplantation for Animal #3 were presented. Left panels: Lentiviral barcodes, Right panels: HDR barcodes.

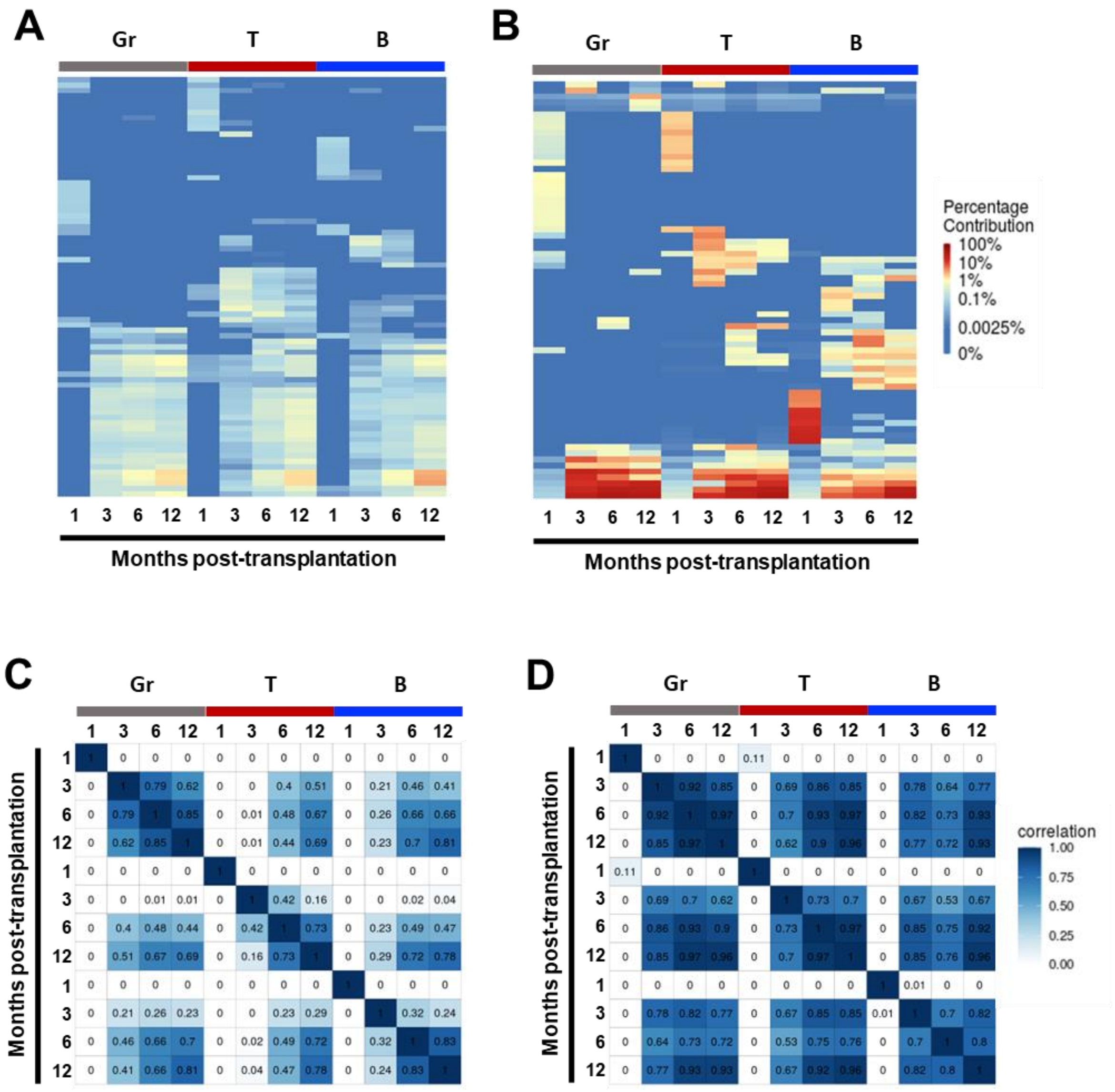


**Figure S7. Clonal contribution analyses across blood lineages in RM *Lentiviral-CRISPR/HDR* competitive Animal #2.** Lentiviral and CRISPR/HDR barcodes were retrieved from purified blood granulocytes (Gr), T cells, and B cells over time for Animals #2. **(A-B)** Heatmaps of the top 10 contributing clones at each time point mapped over all time points. Each column shows a single sample at the indicated time point and each row correponds to an individual barcode (clone) over time. The color gradient represent the relative percent contribution of the barcoded clone in that sample (column) as shown in the color bar on the right. **(A)** Lentiviral barcodes. **(B)** HDR barcodes. **(C-D)** Clonal relationships at a population level as depicted by

Pearson correlation coefficients comparing pairwise fractional contributions from all valid barcodes for the samples across the blood lineage cells at each indicated time point. *r*-values were presented in the each box. **(C)** Lentiviral barcodes. **(D)** HDR barcodes.

**A**

| | DNA | Chromosome | Position | Direction |
|---|---|---|---|---|
| On-target | ATAACGGTTCATACTTCTTTCGG | chr19 | 51026076 | + |
| OT1 | AgAACGGTTCATAtTTCTTTGGG | chr3 | 127176812 | + |
| OT2 | ATAtCGGTcCATACcTCTTTAGG | chr3 | 26154847 | - |
| OT3 | AgAACGaTTtATACTTCTTTGGG | chr3 | 47501751 | + |
| OT4 | ATAACatTTCATAaTTCTTTAGG | chr4 | 79445941 | - |
| OT5 | ATAtCaGTTCATAaTTCTTTTGG | chr6 | 95961899 | - |
| OT6 | AgAACGaTTtATACTTCTTTGGG | chr6 | 168918393 | + |
| OT7 | ATAAaGGTTaATACTTaTTTTGG | chr10 | 89112772 | + |
| OT8 | ActAaGGTTCATACTTCTTTAGG | chr14 | 31223148 | + |
| OT9 | ATgAgGGTTCATAtTTCTTTAGG | chr16 | 76298638 | - |
| OT10 | AgAACGGTgCAaACTTCTTTTGG | chrX | 29807281 | + |
| OT11 | ATAACTGGTTCATACTTtTTTAGG | chr16 | 33580637 | - |

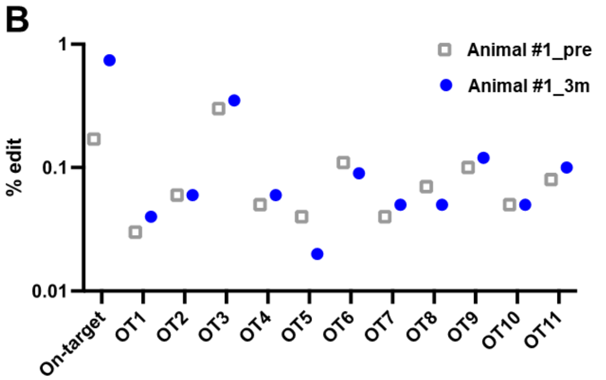


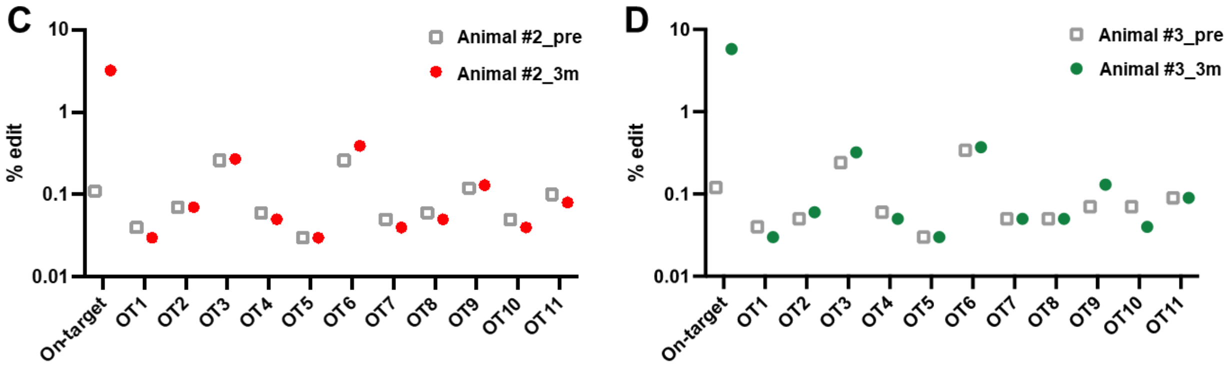


**Figure S8. Off-target analyses for the *CD33* gRNA. (A)** The top predicted off-target sites for the *CD33* gRNA are listed. **(B-D)** DNA from pre-transplantation Gr from each animal was used as negative control (marked as "pre"). Grs were collected from animals' PB at 3 months post-transplantation. Actual off-target events for *CD33* were analyzed by targeted deep sequencing in Animals #1 **(B)**, #2 **(C)**, and #3 **(D)**.

### Table S1. List of candidate gRNAs and DNA oligonucleotides sequence

| # | sgRNA | PAM (NGG) | position | direction | Size (bp) | Primer (FW) | Primer (RV) |
|---|---|---|---|---|---|---|---|
| 1 | GACTCTTGGATCCATAGCCA | GGG | E2 | RV | 20 | CTCCTCCCCAGCTTCCTATC | GTGTGGTAGGGTACGGGATG |
| 2 | GAACACCTCCAATCTTCTCC | TGG | E3 | FW | 20 | GACGCTCCCTCTTGATTCTG | TGAGCTGGATGGTTCTCTCC |
| 3 | TGGTACTGGGAGGGGTTTAG | GGG | i56 | FW | 20 | TCTTTCGGATGGAGAAAGGA | TGGAGTGGTCAGGGTCTAGG |
| 4 | GCAGGAGTCAGTGACAGTAC | AGG | E2 | FW | 20 | AACCTCAGCCTCAGACATGC | TTTGTGGCCACTGGAGAGTC |
| 5 | ATAACGGTTCATACTTCTTT | CGG | E2 | FW | 20 | CAGTGGCCACAAACAAGCTA | CCTAAACCCCTCCCAGTACC |
| 6 | CCTGGTACTGGGAGGGGTTT | AGG | i56 | FW | 20 | TCTTTCGGATGGAGAAAGGA | TGGAGTGGTCAGGGTCTAGG |
| 7 | CAAGTCTAGTGAGAAGAAGC | AGG | i56E3 | RV | 20 | GGTACTGGGAGGGGTTTAGG | ATGAGCACCGAGGAGTGAGT |
| 8 | TGGTCAGGGTCTAGGGCTCC | AGG | E3 | RV | 20 | GGTACTGGGAGGGGTTTAGG | ATGAGCACCGAGGAGTGAGT |
| 9 | CGGAACCAGTAACCATGAAC | TGG | E2 | RV | 20 | ATCCAAGAGTCAGGCTGGAA | TCCTTTCTCCATCCGAAAGA |
| 10 | TACTGGGATCCCCAAGGAGG | CGG | E2 | RV | 20 | GCACTTTCTTCCATCCCGTA | CATGCACAGAGAGCTGGGTA |
| Original | GTCAGTGACAGTACAGGA | GGG | E3 | FW | 18 | AACCTCAGCCTCAGACATGC | TTTGTGGCCACTGGAGAGTC |

| | |
|---|---|
| ssODN | GGCCGATTCCGCCTCCTTGGGGATCCCAGTAGGAACAACTGCTCCCTGAGCATCGTAGATGCCAGGAGGAGGGATAACGGTTCATACTTCGAGC TCNNNNNNNNNNNNNNTTTCGGATGGAGAAAGGAAGTACCAAATACAGTTACAAATCTACCCAGCTCTCTGTGCATGTGACAGGTGAGGCACAGG CTCCAGAAGTG |
| i53 | GCCGCCAGTTTAAACGGCGCGCCATTAATTAAGGATCCAATGTTGATTTTCGTGAAAACCCTTACCGGGAAAACCATCACCCTCGAGGTTGAACC CTCGGATACGATAGAAAATGTAAAGGCCAAGATCCAGGATAAGGAAGGAATTCCTCCTGATCAGCAGAGACTGGCCTTTGCTGGCAAATCGCTGG AAGATGGACGTACTTTGTCTGACTACAATATTCTAAAGGACTCTAAACTTCATCCTCTGTTGAGACTTCGTTGA |

#### Table S2. Primer Sequences for barcode retrieval, targeted deep sequencing, and off-target analysis

##### Primers for barcode retrieval

| Primer name | Sequence (5’----------3’) |
|---|---|
| CD33 CRISPR barcode index forward primer | AATGATACGGCGACCACCGAGATCTACACNNNNNNNNNNNNACACTCTTTCCCTACACGACGCTCTTCCGATCTACGGTTCATACTTCGAGCTC |
| CD33 CRISPR barcode universal reverse primer | CAAGCAGAAGACGGCATACGAGATATCACGGTGACTGGAGTTCAGACGTGTGCTCTTCCGATCTACTGTATTTGGTACTTCCTTTCTCCA |
| Lentiviral barcode index forward primer | AATGATACGGCGACCACCGAGATCTACACNNNNNNNNNNNNACACTCTTTCCCTACACGACGCTCTTCCGATCT |
| Lentiviral barcode universal reverse primer | CAAGCAGAAGACGGCATACGAGATCGTGATACGGCATACGAGCTCTTCCGATCT |

##### Primers for off-target analysis

| Target sites | Forward | Reverse |
|---|---|---|
| OT1 | TGCATGTGTGTCTCTTTATGG | AGGGCCATTTGGTGATTTTT |
| OT2 | TGGGCTGATTCCATATCTTTG | CAACCCAGCAGTCCCATTAC |
| OT3 | TGATATGGCAGCCTCTTTCC | GGTAGCATAAAAGCTTAAAACCTC |
| OT4 | TCAGTAGTGCTGCAACAAACG | AAACACAACTGCCATTTGACC |
| OT5 | GAAGTAGAACCTACCTGTGGATAAA | CGTAAGTTTTATGTCTATGTCGGCTA |
| OT6 | CCTCCCTATCCAAGCACAAG | AAGCCAATAGGCATCACAAA |
| OT7 | TCGGCTATTTCATCCTCCTC | TGCTTGGTCCTTGGAATAGTG |
| OT8 | CCCCACACTGTTTCCCCTAT | ATGGCATGGGATTCTGAGAC |
| OT9 | GATACCACTCCACACCCACTG | TATTTTCCCATCAGCATGGA |
| OT10 | ATGTGCCCCATCCTTGTTAG | TGGTCAAGGAAGTCAAATGG |
| OT11 | GTTGACCTGTGGTTTCAACG | TTGCCCTCCACACTCAAAAT |
